# Power analysis for spatial omics

**DOI:** 10.1101/2022.01.26.477748

**Authors:** Ethan A. G. Baker, Denis Schapiro, Bianca Dumitrascu, Sanja Vickovic, Aviv Regev

## Abstract

As spatially-resolved multiplex profiling of RNA and proteins becomes more prominent, it is increasingly important to understand the statistical power available to test specific hypotheses when designing and interpreting such experiments. Ideally, it would be possible to create an oracle that predicts sampling requirements for generalized spatial experiments. However, the unknown number of relevant spatial features and the complexity of spatial data analysis makes this challenging. Here, we enumerate multiple parameters of interest that should be considered in the design of a properly powered spatial omics. We introduce a method for tunable *in silico* tissue generation, and use it with spatial profiling datasets to construct an exploratory computational framework for single cell spatial power analysis. Finally, we demonstrate that our framework can be applied across diverse spatial data modalities and tissues of interest.

---

Tissues are composed of organized communities of diverse cell types, each with distinct morphologies, molecular profiles and cellular neighborhoods. In homeostasis, cells interact to establish and maintain proper tissue function, whereas diseases can disrupt spatial organization in specific ways^1^. Analyzing such patterns is a cornerstone of histopathology, providing a critical means for diagnosis in disease, and a key tool for understanding tissue function. Molecular measurements *in situ*, especially of RNA and protein markers, enhance the available patterns and aid in mechanistic interpretation. In recent years, emerging methods, including novel spatial transcriptomics and antibody-based spatial proteomics, have dramatically increased the number of molecules that can be measured in one tissue section (**Supplementary Table 1**). This vastly increased the number of possible markers, and in some cases, allowed discovery of new biomarkers *post hoc*^1–3^ in both basic and translational settings^1, 3–6^.

Spatial profiling studies can tackle different key questions, including the association of a specific condition or disease state with particular cell types, cell-cell interactions, or higher-order structures in the tissue. To address such questions, scientists need to design experiments, including choosing the number of samples and the number and size of fields of view (FOVs) required to detect spatial patterns at a given confidence level. Each of these choices depends on specific assumptions, such as the organization of the tissue, the type of measurements, variation within and between specimens (and classes), and the statistical methods applied.

To the best of our knowledge, statistical frameworks tailored for such power analysis for spatial profiling methods are currently lacking. Prior power analysis methods in genomics were devised in the context of either bulk profiling, where the tissue is homogenized, or single cell profiling^7–12^, where cells are dissociated (**Fig. 1a**). In suspension experiments, there are few relevant parameters for sampling strategy - the overall number of cells and the relative abundance of different cell types (**Fig. 1a).** So far, spatial profiling studies have focused on detecting spatially-resolved genes or specific cellular neighborhoods *post hoc*^13–16^, but did not consider questions of sampling strategy, such as the number of specimens or FOVs needed to reliably detect different patterns, nor the effect of FOV size (**Fig. 1b**). Finally, power analyses previously performed to address heterogeneity of single (bio)markers in whole tissues do not scale to novel profiling technologies^17^.

**Fig. 1:**
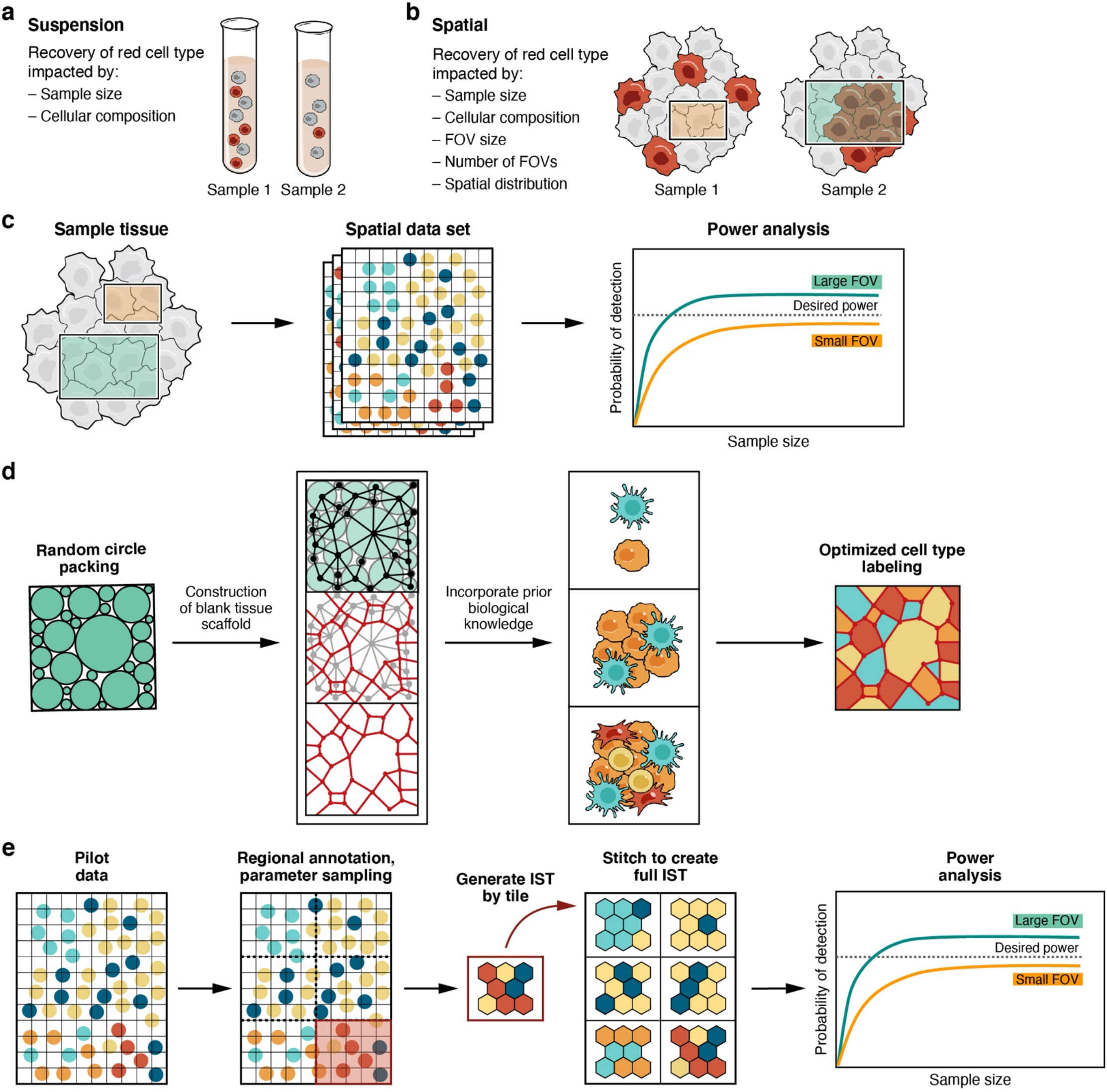
Power analysis framework for spatial omics data. **(a,b)** Features impacting power to detect cell types in single cell and spatial genomics experiments. **(c)** Use of spatial data sets for retrospective power analysis. Different sizes of FOVs (squares, left) from an existing spatial dataset are sampled, and their data (middle) is used to conduct a statistical analysis. The results (right) are used to calculate the probability of detection of a desired feature (y axis, right) when using smaller (orange) or larger (green) FOVs. Dashed line: desired threshold. **(d)** Generation of ISTs. From left: Our method generates a blank tissue scaffold using a random circle packing algorithm (two left panels), and prior biological knowledge is used to optimize cell type assignments on the tissue scaffold (second from right), followed by visualization with Voronoi diagrams (right). **(e)** IST generation of complex or large tissues by regional annotations from pilot data. Pilot data (left) is used to assign regional annotations (second left), and spatial parameters are estimated for each region separately. The region-specific parameters are used to generate IST tiles, which are stitched together to create a full IST (second from right), followed by analysis, for example to compare the sampling requirements to detect a spatial feature at a desired power (dashed line) for a small (orange) vs. large (green) FOV.

Spatial power analysis poses several challenges. First, spatial experiments offer a very large number of possible spatial features that might be relevant, and these features may be challenging to pre-define. Thus, in addition to distribution of cell type proportions (as in single cell genomics), cellular organization in the context of other cells and the tissue architecture are paramount, but such structures are difficult to parameterize and vary across tissues. Second, power analysis usually requires exploration of large amounts of data or a well-defined model of the system of interest to simulate the underlying distributions. While in some settings (e.g., addressing how field of view (FOV) size impacts feature detection in one slide) it is possible to proceed directly from limited spatial data to power analysis (**Fig 1c**), other questions (*e.g.*, how many whole slide images are required to detect all significant cell-cell interactions in a cohort) requires substantially more data, which may not be available at this time.

To begin to address these challenges, we introduce a power analysis framework to help design and interpret spatial profiling studies in tissues, including an approach to generate tissues *in silico* by parameterized models of tissue structure, overcoming limited data availability and serving as an approximate generative model for tissues. First, we construct blank tissue structures (“tissue scaffolds”) and apply heuristic and optimization-based labeling solutions to generate *in silico* tissues (ISTs) that reflect parameterized spatial features and molecular information (**Fig. 1d, Methods**). To generate a tissue scaffold, which represents the spatial location of generic cells, we employ a random circle packing algorithm to generate a planar graph. Next, we assign an attribute labeling to the graph, where attributes on nodes represent cell type assignments. The labeling is based on two data-driven parameters for a given tissue type: the proportions of the *k* unique cell types, and the pairwise probabilities of each possible cell type pair being adjacent (a *k* x *k* matrix) (**Fig. 1d, Supplementary Fig. 1, Methods**). We assume that these data-driven input parameters are available from prior knowledge or a pilot phase of a study. These parameters are local in nature and could vary across the tissue. For instance, samples with known gross morphological regions may have different cell type abundances and adjacency probabilities in each region. In such a case, using prior knowledge of the gross morphology, we generate sub-regions drawn from parameters corresponding to morphological regions, and stitch them to create a full IST (**Fig. 1e**, **Methods**). This generates a mosaic representation of tissue architecture. We then use this feature-independent framework to directly perform and validate power analysis results. Note, that while we used cell type labels as attributes, any type of attribute can be used.

First, we used ISTs for experimental design focused on cell type discovery in spatially-resolved data, considering two sampling strategies: one where single cells are observed in isolation from their spatial context, analogous to (non-spatial) single cell profiling methods, and another when spatially contiguous fields of view (FOVs) are observed (**Fig. 1a,b**). We constructed two statistical models to describe the corresponding probability of cell type discovery in spatial sampling: a beta-binomial model to predict how many single cells need to be measured to observe a cell type of interest at a certain probability and a gamma-Poisson model to predict how many FOVs are required to observe a cell type of interest at a certain probability (**Methods**). We then applied our framework to demonstrate how ISTs can be used to help experimental design for cell type discovery in spatial profiling experiments. As a case study, we generated small ISTs with 2,186 cells, which approximates 500×500 μm, a typical size of one core in a tissue microarray (TMA)^1,5^. Next, we labeled cells with 4 different cell types in three spatial configurations: (**1**) a tissue where a rare cell (3% abundance) is randomly located (**Supplementary Fig. 2a**, maroon); (2) a tissue with one cell type exhibiting strong self-preference (**Supplementary Fig. 2b,**purple), and (3) unstructured tissues (to serve as a null model), where cells of all types have equal probabilities of being adjacent to any other cell (given their proportions) (**Supplementary Fig. 2c**).

As expected, cell type abundance greatly affected the number of cells and FOVs required to have a specified likelihood of observing a cell type of interest. For example, after sampling 20 cells in our null tissue, we are nearly guaranteed to observe a common cell type of interest (abundance 22%) at least once, whereas for a rare (3%) cell type of interest, sampling 100 cells gives just an 80% chance of discovery (**Supplementary Fig. 3a**). Moreover, for ISTs with the rare cell type design, we asked how many FOVs of a fixed size (1%, 5%, 10% of tissue area) are required for a given probability of observing the rare cell type in at least one FOV (**Supplementary Fig. 3b**). For example, at least 3 FOVs each of 1% of the tissue size (~22 cells) must be examined for observing the rare cell type in at least one FOV at 80% probability (**Supplementary Fig. 3b**).

We then used ISTs to determine the sampling strategy required to detect cell-cell interaction patterns in a set of samples as compared to a null model. To this end, we applied a permutation test^5,18^ to identify pairs of cell types that occur in proximity more (“significant interactions”) or less (“significant avoidances”) frequently than expected by chance (**Methods**), by comparing a real tissue to a null model. To simulate this setting, we generated two sets each of 25 ISTs (2,186 cells per IST) – one structured by self-preference of one cell type (to simulate real tissue) and another following a random tissue model (to form the null) – and identified cell-cell interactions that characterized structured ISTs compared to the random (null) tissue model (permutation test p<0.01, **Methods**). Hierarchical clustering of the permutation test results showed that the self-preference ISTs consistently had the desired interaction, but the randomly structured set did not (**Supplementary Fig. 4a**). To simulate a more complex structure, we generated another set of 25 structured ISTs now with an enriched interaction between three of 10 cell types, and 25 random ISTs with the same 10 cell types but without any constraints on the interactions. Again, hierarchical clustering of the permutation test scores for each pair of cell types separated structured ISTs from non-structured ISTs, with the enriched interaction recovered only in the structured set (**Supplementary Fig. 4b**). Next, we showed how tissue sets were separable based on interactions. Therefore, we tested whether the distributions of interaction significance scores for each interaction were significantly different between the structured and unstructured ISTs for different numbers of tissues. We found that the specified interactions were among the interactions with distinguishable score distribution even when only a small number of tissues were compared (**Supplementary Fig. 4c**).

Next, we applied our approach to parameters derived based on three real biological datasets: a high-density spatial transcriptomics (HDST) dataset of breast cancer, an osmFISH dataset of the mouse cortex, and a highly multiplexed antibody-based (CODEX) murine spleen dataset (**Supplementary Table 1)**. In each case, we used available gross morphological data to estimate cell type abundance and pairwise adjacency probabilities from each annotated morphological region in the dataset and generated ISTs on a tile-by-tile basis using region-specific estimates of spatial parameters (**Fig. 2a-l**; **Methods**). We selected these datasets because they span a broad range of complexity of biological structure. The HDST breast cancer data is relatively unstructured; despite provided annotations of morphological zones, the tissue is dominated by one cell type (epithelial cells) with little variation in composition between morphological zones (**Fig. 2a-b, Supplementary Fig. 5**); the mouse cortex is a highly ordered, layered tissue with unique cell types in each morphological zone (**Fig. 2e-f**); and the mouse spleen has complex, recurrent structure with shared features between morphological zones of the same type (**Fig. 2i-j**).

**Fig. 2.**
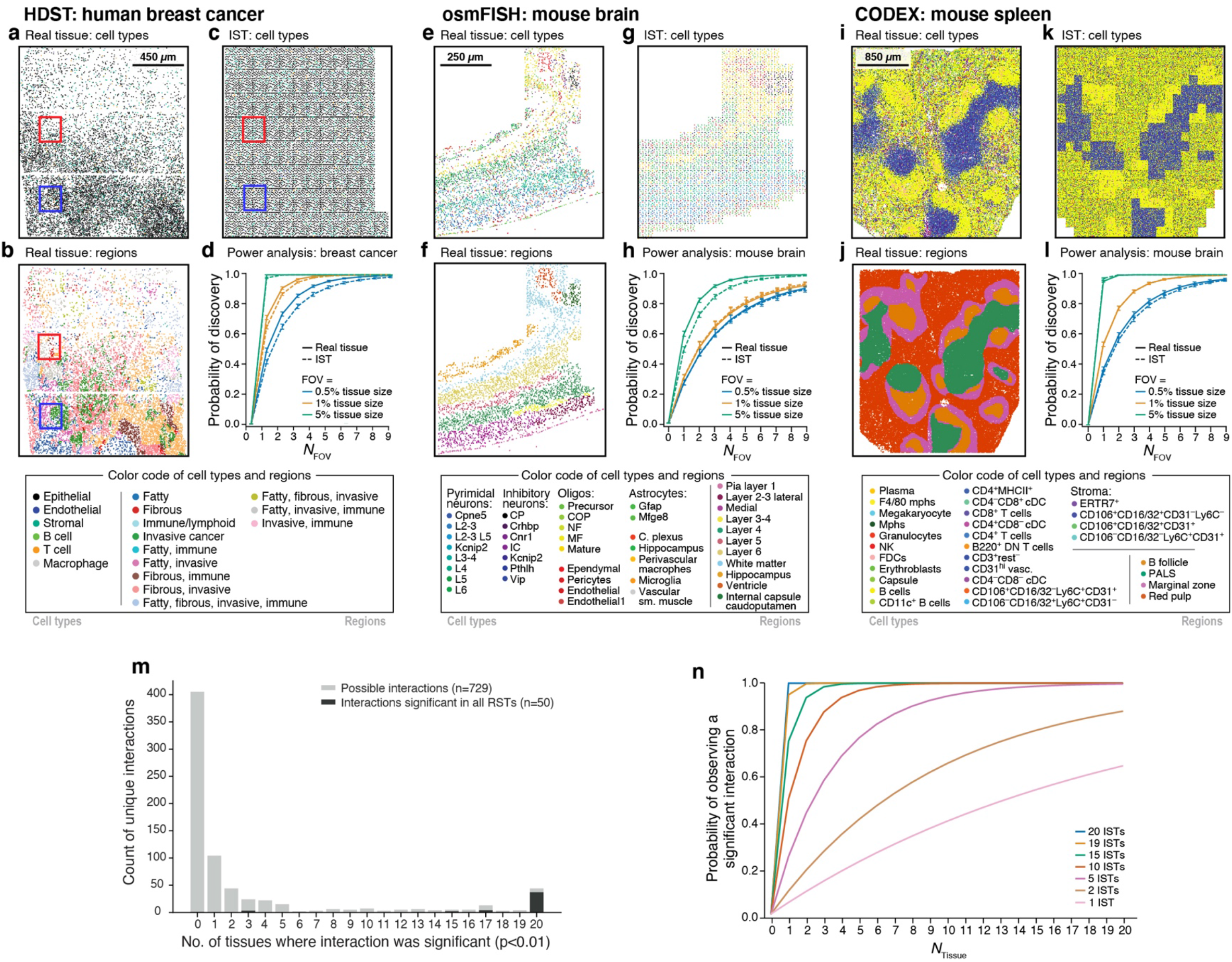
Spatial power analysis to recover cells and cellular interactions of interest using ISTs in different tissue types. (**a-l**) Power analysis for number and size of FOVs required to detect rare cell type by spatial analysis of tissues of different structure. **(a-d)**HDST dataset of breast cancer. **(a-c)** Cells (points) at their spatial position in real (a,b) and corresponding IST (c) data, labeled by type (a) or morphological regions (b). Red and blue rectangles are expanded in **Supplementary Fig. 5**. **(d)** Probability (y-axis) to discover at least one T cell when sampling different number of FOVs (x-axis) of different sizes (colored lines), in either real tissues (solid lines) or ISTs (dashed lines). **(e-h)** osmFISH murine cortex data. (**e-g**) Cells (points) at their spatial position in real (e,f) and corresponding IST (g) data, labeled by type (e) or morphological regions (f). **(h)** Probability (y-axis) to discover at least one L6 Pyramidal neuron when sampling different number of FOVs (x-axis) of different sizes (colored lines), in either real tissues (solid lines) or ISTs (dashed lines). **(i-l)** CODEX mouse spleen data. (**i-k**) Cells (points) at their spatial position in real (e,f) and corresponding IST (g) data, labeled by type (i) or morphological regions (j). **(l)** Probability (y-axis) to discover at least one megakaryocyte when sampling different number of FOVs (x-axis) of different sizes (colored lines), in either real tissues (solid lines) or ISTs (dashed lines). **(m,n)** Power analysis for number of tissues required to detect a significant cell-cell interaction. **(m)** Distributions of number of unique cell-cell interactions (y-axis) detected as significant (p<0.01, permutation test) in a set of 20 ISTs (x-axis). for 729 possible cell-cell interactions (light grey) and for the 50 cell-cell interactions that were significant in all three real spleen tissues (dark grey). **(n)** Probability of observing a significant interaction (y-axis) for different number of tissues sampled (x-axis) for interactions recovered in different numbers of ISTs (line color).

Power analysis shows how the extent and nature of tissue structure impacts the number of cells and FOVs required for cell type discovery. In each dataset, we selected a lowly abundant cell type to better illustrate the effect of sampling strategy on feature recovery (as highly abundant cell types would be detected universally). We compared two sampling strategies: (**1**) sampling FOVs and assaying them in their entirety (“spatial sampling”) for the presence of a cell type of interest (*e.g*. analysis of a TMA or a specified ROI**; Fig. 2d,h,l**) and (**2**) dissociated single cell analysis (“single cell sampling”) of the *entire* tissue sample, such that no spatial information is retained (*e.g.* flow cytometry or scRNA-seq; **Supplementary Fig. 6b,d,f**). For spatial sampling, we used a gamma-Poisson model to determine the number of FOVs of a fixed tissue area (FOVs may have varying cell counts due to different cell densities, which is accommodated by the model) required to detect a cell type of interest in real tissue or in its corresponding IST (**Fig. 2d,h,l**). We assumed that a cell type is completely determined by its markers and defined detection as observing at least one cell of the type in an FOV. For non-spatial single cell sampling, which does not capture any spatial information and is equivalent to a FOV sized to capture one cell, we employed a binomial process with the same assumptions.

With spatial sampling, in the relatively unstructured breast cancer tissue, there is an 80% probability to detect a T cell in one FOV of 5% (~500 cells) total tissue size (**Fig. 2d**). In the mouse cortex, where the tissue is highly structured and non-repetitive, discovering the more abundant L6 pyramidal neurons (9% abundance) at the same 80% probability (**Supplementary Fig. 6c**) requires two FOVs of 5% (~650 cells total) of tissue area (**Fig. 2h**). Finally, in the mouse spleen, the repeating morphological structures (*e.g.*, periarterial lymphatic sheaths (PALS) and B follicles surrounded by a marginal zone) lower the number of FOVs required to recover even very rare cell types like megakaryocytes (~0.1% abundance): just one 5% FOV (~4300 cells) is sufficient to detect at least one cell at >80% probability (**Fig. 2i-l**). A megakaryocyte can also be captured at 80% probability by sampling 4 FOVs each at 0.5% tissue area (~1700 cells total), illustrating the impact that sampling strategy has on the absolute number of cells required to detect a spatially distributed feature. Smaller FOVs are less impacted by overdispersion (and when sized as one cell, are equivalent to single cell sampling). When a cell is overdispersed in the context of a highly ordered and heterogeneous tissue (mouse cortex), multiple smaller FOVs yield better detection probability than a single larger FOV (**Fig. 2h**), but this is not the case in tissues with more repeatable organization (spleen). While we normalized sample size, absolute tissue size is important because biological features exist at different length scales (*e.g.*, a sample entirely within a tissue sub-region that lacks a cell type will never result in the discovery of that specific cell type). By comparison, with non-spatial single cell sampling, in breast cancer, profiling ~100 cells would achieve an 80% probability of detecting at least one T cell (**Supplementary Fig. 6a,b**), 17 cells suffice to detect at least one L6 pyramidal neuron at 80% probability in the mouse cortex (**Supplementary Fig. 6c,d**), and ~1,270 cells are required to detect a rare megakaryocyte at 80% probability in the mouse spleen (**Supplementary Fig. 6e-f**). Thus, power analysis considering only overall cell frequencies would vastly underestimate the FOVs required for a spatial experiment.

Next, we used our framework to detect significant cell-cell interactions in real data. We defined significant interactions and avoidances via a permutation test^5,18^, as described above (**Methods**), determined the number of FOVs required to detect any significant finding, and estimated how FOV size selection impacts the types of detectable interactions. Focusing on spleen as a case study, we examined CD4^+^ and CD8^+^ T cell interactions, which are enriched in the full tissue (p<0.01, permutation test, **Methods**). Using our IST, we estimate that measuring >7.5% of the assayed tissue size (~123×123μm, ~5,600 cells) would recover this interaction as significant (permutation test, p<0.01) at 80% probability, with a sharp inflection point (**Supplementary Fig. 7a,b**). This inflection point should be accounted for when sampling with fixed FOV sizes, as in the case of tissue microarrays (TMAs); TMAs of insufficient size may never capture the feature of interest. In general, areas in which the interaction of interest is recovered span across morphological zones, such that they are representative of the diversity of tissue structures (**Supplementary Fig. 7c**, green squares).

In certain experimental designs, researchers may desire to ask whether the enrichment of a specific significant interaction is different between two tissues (or two subsets of tissues) and assess sampling requirements to achieve statistical power to detect differentially significant interactions. To assess this in the context of CD4^+^ and CD8^+^ T cell interactions, we analyzed the real mouse spleen dataset along with a copy where we rearranged cells in adjacent CD4^+^ / CD8^+^ pairs to reduce the CD4^+^ and CD8^+^ T cell interaction frequency by 37%, while preserving the overall cell type frequency and tissue structure (**Methods**). Next, we quantified the enrichment of a specific cell-cell interaction by the Interaction Enrichment Statistic (IES), which we defined here as the frequency of a specific interaction relative to the frequency expected given the proportion of the two cell types (**Methods**). We then drew 100 FOVs of fixed size (5%, 7.5%, 10% of full tissue size) from each of the two tissues and calculated the IES in each FOV, yielding an IES distribution. Finally, using the maximum likelihood estimate of the mean and variance, we fit a Gaussian to the IES distribution.

FOV size has a substantial impact on the ability to detect differentially significant interactions between samples (**Supplementary Fig. 8**). With a 5% FOV, the CD4^+^ and CD8^+^ T cell interaction is significantly detected only rarely (**Supplementary Fig. 8a**), and we cannot identify it as differentially significant between the two tissues (p=0.41, Z-test; **Supplementary Fig. 8a**). When the FOV size is increased, the differentially significant interaction is readily detected (7.5% FOV, Z-test p=0.018; 10% FOV, Z-test p<<0.01). Finally, we systematically tested how sample size and effect size affect power (**Supplementary Fig. 8d-f).** Because IES measures enrichment relative to the proportions of cell types present in a sample, it assumes that these proportions are equal between samples.

Finally, we showed how our *in silico* framework can be used to make predictions of sampling requirements when the set of true features of interest is unknown (in contrast to pre-specified cells or interactions above). To this end, we assembled a set of three real mouse spleen tissues, estimated the input parameters for IST generation from one of these three tissues, and held the remaining two for validation. We generated 20 ISTs based on the estimated parameters; a number selected to capture a broad set of cell-cell interactions that can be spuriously detected as significant given the input parameters or biological noise (in real data). Unlike in previous analyses, we aimed to enumerate a set of statistically significant spatial features, rather than recover a known ground truth. Given this goal, and the fact that our tissue generation approach does not recapitulate macrostructures natively, there is a risk that repeating macrostructure layout in all ISTs could generate spurious interactions. To address this, we shuffled macrostructures based on regional annotations included in the real dataset (**Methods**), and then called significant cell-cell interactions in the ISTs individually and in the dataset overall (Permutation test, p<0.01). Of 729 possible pairwise interactions, only 69 were significant in more than 80% of ISTs, of which 44 were significantly enriched in all 20 ISTs (**Fig. 2m,**grey). Importantly, of the 50 interactions that were significant in all three real tissues, 37 (84%) overlapped with the 44 significant in all 20 ISTs (**Fig. 2m,** black). Another 13 were idengified as significant in real spleen data but were not among the 44 interactions that were detected as significant in all ISTs and were largely associated with cell types at the boundaries in the segmentation mask or tissue (**Supplementary Fig. 9b**). To predict the number of samples required to observe a specific interaction at a desired probability, we calculated the proportion of ISTs in the set in which we observed a specific interaction (**Fig. 2n**). For example, to detect at 80% probability an interaction of interest that occurs in just 5 of 20 ISTs, an experiment should have at least 6 tissue samples.

In conclusion, we developed an *in silico* tissue framework to enable spatial power analysis and assist with experimental design. ISTs can be directly used for method development and benchmarking of existing^15,16,18,19^ or novel spatial analysis methods. Our framework is agnostic to feature type and assigned labels can be derived from any type of information assayed in the tissue. Here, we used cell type labels instead of individual quantitative features (*e.g*., marker intensity or cellular morphology) to provide a straightforward and interpretable abstraction, but any spatial profiling data can be used. In all cases, our power analysis based on individual ISTs accurately predicted the probability of cell detection compared to the real tissue, showing that IST generation has mimicked actual tissue structure given estimated parameters from a variety of spatial profiling data types and underlying tissue structures (**Fig. 2a-l**). Overall, we robustly created ISTs across diverse tissue types and various experimental methods to perform accurate spatial power analysis for cell type detection. While retrospective power analyses could be performed on sufficiently large extant biological datasets, this is not necessarily practical for designing new spatial experiments, where the particulars of spatial structure impact power. As an alternative, ISTs enable predictive spatial power analysis to inform experimental design decisions early in a study, depending on the feature of interest. We provide a tool to create ISTs, perform statistical testing to identify spatial features, simulate different experimental design choices and perform spatial power analysis. Using this framework, we enumerated some parameters for consideration in the design of spatial experiments, including tissue size, diversity of cell types, spatial structure, sampling strategy (*e.g.*, TMA size selection), and feature of interest (*e.g.*, cell type discovery, spatial motif discovery). Our work will help towards extracting meaningful biological or clinical insights from spatial genomics studies.

## Methods

### *In silico* tissue generation

To perform spatial power analyses, we devised an approach to simulate tissue *in silico*. *In silico* tissues were generated by first constructing a tissue scaffold - a blank tissue with no cell information assigned - then assigning cell type labels to the scaffold.

### Generating tissue scaffold

Tissue scaffolds were generated with a random circle packing algorithm. This algorithm places circles of a bounded random radius within a rectangular region, disallowing overlaps between circles via rejection sampling. The algorithm continues until it fails to place any new circle 500 consecutive times. This results in a densely packed region. In this model, circles represent cells. Touching circles represent adjacent cells and will be connected by an edge in the graph representation (**Fig. 1d**).

Circle packing results are then converted into a graph representation. A graph is a highly interpretable data structure that can represent a tissue due to its clear encoding of spatial relationships and ability to be labeled with biological information. This is performed by calculating, for each circle, all other circles within the smallest allowable radius of the original circle's perimeter. Effectively, for a circle *C*, this finds all circles that *C* would overlap with if the radius of *C*, *r_c_* was modified such that *r_c_* = *r_c_* + *r_min_*, where *r_min_* is the smallest radius. These circles are considered to be adjacent to *C*. A node is placed at the center of each circle and an undirected edge is drawn to the node corresponding to each of the adjacent circles (**Fig. 1d**).

### Assigning cellular information

After generation of the tissue scaffold, cellular information was assigned to the tissue. We specified two input parameters in this process, a vector *p* ∈ ℝ^*k*^, which contains the probabilities of discovering each of the *K* cell types in the tissue. Further, we define a matrix *H* ∈ ℝ^*k* × *k*^ where *h_ij_* ∈ *H* defines the probability that a cell of type *k_i_* is adjacent to a cell of type *k_j_*. Two algorithms were used to assign labels to the tissue scaffold.

### Graph neighborhoods/heuristic assignment

A neighborhood, *N*_υ_, was defined on the graph, *G*, representation of the tissue scaffold. For a vertex *υ* ∈ *G*, *N_υ_* = *G*[*S*] is defined as the subgraph induced by the set *S* = {*u* ∈ *G*|*d*(*v*, *u*) ≤ *∈*}, where *d* is a function computing the geodesic distance and *ϵ* specifies the search radius.

The graph region was partitioned into a grid of *50* × *50* px regions. Within each region, a start node *v*_i_ was selected at random. The type 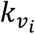 of *v*_i_ was sampled from a multinomial distribution of the cell type probabilities: 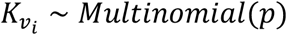. Given the choice of *k*, the probabilities of the type labels for the nodes 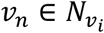 are sampled from a multinomial distribution of the corresponding row vector in *H*, *v*_n_ ∼ *Multinomial*(*H*_k∗_).

The partition grid then is shifted horizontally and vertically by 25 px, and the sampling process repeated. Any remaining unlabeled nodes are then discovered and assigned by the same process. After all nodes are labeled, random nodes are selected, and the observed neighborhood label distribution 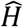 is calculated and compared to *H*_k∗_. Overabundant type labels in 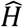 are swapped to under-abundant type labels.

### Optimization of cell assignment

### Setup

Given the blank tissue scaffold, we compute an assignment matrix *B* ∈ ℝ^*n* × *k*^ that describes the cell type assignment for *n* cells in the tissue scaffold. An entry *B_vk_* = *1* if a node *v* is of type *k*, else *B_vk_* = *0*. Furthermore, as each cell may only receive one type assignment, each row in *B* sums to 1, 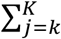 *b_ij_* = 1 and each column sums to the expected cell type count, which, when normalized, yields the cell type distribution *p*. In a fully labeled tissue with adjacency matrix *A*, the matrix of neighborhood probabilities given an assignment *B* can be computed as 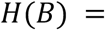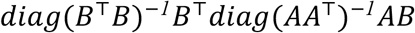.

### Objective

Given a target matrix of neighborhood probabilities *H*(*B*^∗^) derived from real data and a random synthetic tissue scaffold with its resulting adjacency matrix *A*, we aim to generate probabilistic synthetic assignments of cells to labels that conserve observed neighbourhoods of cell label to cell label preferences.

We formulate this problem as an inverse optimization problem, in which we seek to find a probabilistic assignment matrix *B* ∈ ℝ^*n* × *k*^ that would lead to a matrix of neighborhood probabilities *H*(*B*) matching the observed data as closely as possible.

The resulting objective aims to recover a matrix *B* representing synthetic data that optimizes the loss

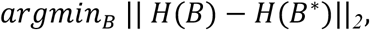

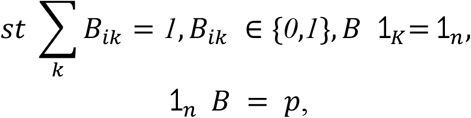

where, as before, *p* is the cell type distribution we aim to match and where ⥠_*k*_ represent a *K* dimensional vector of ones. When the assignment is required to be unique and all the entries of *B* are integers, the question of whether such a labeling exists is generally difficult to settle. In a particular case, if cells sharing the same label exhibit strong repulsive behavior towards one another such that the neighborhood probabilities *H*(*B*^∗^) is a matrix with zero diagonal, without the constraint ⥠_*n*_ *B* = *p*, the optimization problem is akin to the well known vertex graph coloring problem^20^. In the *k*-coloring vertex problem one wants to decide whether a graph can be colored using *k* colors such that no vertices of the same color share an edge. For *k* > *2* this problem and many of its variants are known to be NP-complete.

The considered loss is further equivalent to the semidefinite program objective

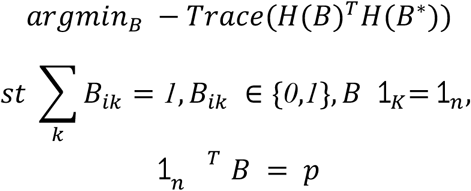

Finally, we derive an efficient algorithm to solve a relaxed version of this problem by considering the augmented Lagrangian objective over a matrix B with continuous entries

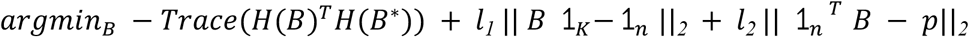

for positive real parameters *l*_*1*_ and *l*_*2*_.

### Implementation details

For GPU accelerated automatic differentiation, we implemented the optimization routine using JAX in Python 3.7 (http://www.github.com/google/jax). We provide further details regarding implementation, system requirements and demo instructions online at https://github.com/klarman-cell-observatory/PowerAnalysisForSpatialOmics. For details regarding optimizing the augmented Lagrangian objective, see the function optimize_assignment in the spatialpower.tissue_generation.assign_labels module.

### Parameter optimization

The user directly supplies the expected cell type proportion, *p*, and the expected neighborhood distribution matrix, *H*. The optimization routine has additional parameters. We set the learning rate as well as two additional loss weight parameters, ‘l1’ and ‘l2’. The two parameters ‘l1’ and ‘l2’ weigh the relative contribution of constraints on the bounds of the probabilistic assignment and *p*, respectively. In detail, the first parameter enforces that all the n rows of *B* sum to one, while the second one enforces that the resulting solution *B* marginally matches cell type proportions (columns sum to the desired expected numbers of cells of a given cell type).

Note that in its current form, the objective enforces (through the term dominated by ‘l2’) that the assignment *B* matches the cell type proportions uniformly. Since the constraint parameters are additive, we can, however, encourage our objective to be more biased toward populations of cell labels which, due to their rarity, might otherwise be overlooked. We can accomplish this by introducing optional, cell label specific parameters *w*_*k*_ to control the relative contribution of the specific constraints on cell label proportions *p*, invoking a tradeoff between unique assignment and matching assignments to *p*. The corresponding objective is

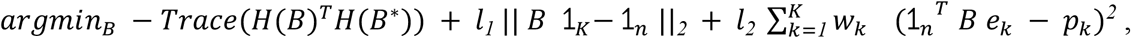

where *e_k_* is the standard basis vector of dimension *K* with non-zero value at index *k*. For example, when dealing with a rare cell type -- low *p_k_* -- a higher weight *w_k_* will enforce that the rare cell type is going to have a nonzero chance of appearing in the resulting synthetic cell assignment.

### Incorporating cell type proportion information

Due to inherent tradeoffs between optimizing with respect to *P* and *H* jointly - our objective is sufficiently close to a graph coloring problem that ideal solutions may not be possible - we attempt to assert control over which specific interactions are favored in the optimization process. Since it seems more likely that a user may have prior knowledge about which interactions are the most or least abundant (“extreme values”) we provide an option to optimize only over those elements of *P* that are beyond one standard deviation from the mean. This favors extreme values in *P* by changing which values in *w_k_* the *l*_*2*_ constraint is applied to (see ‘extreme_values’ in ‘constraint()’ in the tissue_generation.assign_labels.optimize module).

### Cell type heterogeneity

To model the number of cells that must be measured to achieve a desired probability of observing a cell type of interest in a single cell profiling experiment, we calculated the proportion of cells in a tissue that were of the type of interest. We used a simple binomial model to predict the number of cells that need to be profiled to achieve a certain probability of observing the cell type of interest at least once.

We model the number of FOVs with a certain count of cells of a particular type of interest that an experimentalist can expect to observe in an experiment. Due to underlying tissue structure that results in overdispersion in the counts of cell types of interest per FOV, we construct a gamma-Poisson (negative binomial) model. The negative binomial distribution (NBD) can be parametrized in several ways, but here we consider the following parameterization: 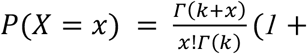 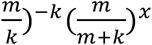 for *x* ∈ ℤ^0+^ and where *m* > *0* and *k* > *0* are parameters describing the mean and shape, respectively. We compared estimating NBD parameters by methods of moments estimation and the zero term method (ZTM)^21^. Due to the high frequency of FOVs with no cells of the type of interest, we found the ZTM estimator achieved superior performance. We estimated 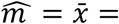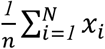. To estimate 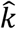, we numerically solve the equation 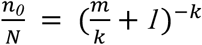 where *N* is the sample size and *n*_*0*_is the count of zeros. The numerical solution was computed with the ‘fsolve’ function in SciPy^22^. A probability of discovery was computed by computing the complement of the model evaluated at the zero count, but the NBD describes the probability of describing any number of cells of the type in the FOV. Furthermore, the NBD can accommodate the fact that FOVs vary in the number of cells they contain (e.g. because of differences in cell density across tissues). Importantly, this model also assumes that a specific combination of makers has 100% accuracy to define the cell type label.

### Full image creation

To construct ISTs at the scale of whole slide images, we can compartmentalize the image into distinct morphological regions representing unique macrostructures (**Fig. 2**). All datasets selected for this analysis contained domain expert macrostructural annotations. For each, we estimated the required parameters for tissue generation from all annotated macrostructural zones. We generated a lower-resolution segmentation map by partitioning the original segmentation map into a grid and determining the dominant zone in each grid partition. We generated small tissue scaffolds based on the mean number of cells per grid partition and generated one assignment solution per grid square (tile). The parameters used in each tile matched the dominant zone in that tile. Tiles were stitched together to generate a composite image (**Fig. 2**). To save computational time in large images, we generated only one blank tissue scaffold and relabeled it for each tile. This approach additionally enables simpler stitching of tiles, though does create an argifact during visualization because of a high density of points on the boundaries. Because our model only considers graph connectivity, this is only a drawback during visualization.

### Visualization

For small ISTs, we generate a tissue-like representation by computing a Voronoi diagram and coloring each Voronoi region with a color representing the cell type assignments **(Supplementary Fig. 2**). For larger ISTs, computing the Voronoi diagram can be slow. In this case, we simply visualize the tissue as a scatter plot, colored by cell type assignment.

### Tile shuffling

To determine the effects of tissue macrostructure in the murine spleen data, we generated 20 full-size ISTs with randomized macrostructure. Each of these ISTs contains the number of tiles from each zone as found in the original segmentation map. Tiles were generated as described above, but randomly stitched to generate shuffled images.

### Neighborhood discovery via permutation testing

We generated tissues with a significant pairwise cell-type interaction, idengified as pairs of cell types that occur both more frequently (“significant interactions”) and less frequently (“avoidances”) than expected via a permutation test^5,18^. In the permutation test, the ground truth neighborhood distribution is calculated. Then, the assignment labels on the tissue are shuffled to relabel the tissue, preserving the tissue structure. At each shuffle, the neighborhood distribution is recalculated. A p-value for each interaction pair is calculated as the fraction of observations that are more extreme than the ground truth value. This test is performed twice, once for each tail of the distribution which provides a directionality for the interaction (e.g. cell type A surrounded by cell type B or vice versa).

### Clustering for motif interaction discovery

We performed agglomerative hierarchical clustering to verify that cohorts with parameterized spatial distributions and spatial null cohorts exhibited the expected significant interactions and avoidances as well as identify motifs of more than 2 cell types^18^. For a given *in silico* tissue, we performed a permutation test for each possible interaction (for *k* cell types, there are *k* × *k* possible interactions) and calculated a p-value. We then clustered based on these scores using the unweighted pair group method with arithmetic mean (UPGMA) algorithm, as implemented in Scipy v. 1.4.1.

### Interaction enrichment statistic (IES) and Z-test

We defined a statistic to quangify the overall enrichment of a cell-cell interaction relative to an expectation based on the proportion of cell types and a linear algebraic method for fast computation. As a theoretical framing, consider a tissue to be an undirected graph *G*(*V*, *E*) in which cells are represented by vertices and an edge represents a direct adjacency between two cells. The *K* types are encoded in the graph as attributes on the vertices. For an interaction between two cells of type A and B, we define the expectation of the number of edges in the graph that connect a cell of type A and a cell of type B as *∑* = *2f_A_f_B_*|*E*| where *f_k_* is the frequency of a cell of type k. Then, we define the IES as 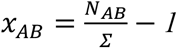, where *N_AB_* is the number of edges connecting a cell of type A with a cell of type B. An IES of 0 indicates no enrichment over expectation, negative and positive values indicate depletion and enrichment, respectively.

To conduct a test of difference between two IES distributions, we calculate 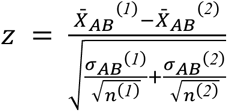 where 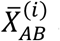 and σ_AB_^(i)^ are the sample mean and standard deviation of IESs between cells of type A and B in sample 1. The probability of z is calculated using a standard Gaussian survival function.

We devised the following method to efficiently calculate the IES in complex graphs. Let *A* be the adjacency matrix corresponding to *G* and *B* be a |*V*| × *K* matrix of one-hot encodings of cell type. Let *i* and *j* be the indices corresponding to the one-hot encoding of types X and Y, respectively. We construct the symmetric matrix *C* = *B*^⫟^*AB*. If *i* ≠ *j*, the element *C_i,j_* is equivalent to the number of edges (*i.e.*, interactions) between types X and Y. If *i* = *j*, the number of edges between two cells of the same type is 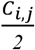.

### Retrospective power analysis

We conducted a retrospective power analysis by generating tissues with three different spatial compositions. Via permutation test, we compiled a list of all significant interactions and avoidances in each generated tissue to establish ground truth of the full diversity of spatial interactions in a sample. Then, we drew contiguous spatial samples of increasing size and conducted a permutation test to identify significant interactions and avoidances within the subsample. We compared the idengified significant interactions and avoidances from the subsample to the ground truth and calculated the proportion of ground truth spatial interactions that were recovered in the subsample and the proportion of falsely called significant interactions and avoidances in the subsample. For each size increment, 100 trials were conducted.

## Acknowledgments

The work described in this article contributed towards the goals of the Human Tumor Atlas Pilot Project (HTAPP: task order no. HHSN261100039 under contract no. HHSN261201500003I, NCI, National Institutes of Health) under the Human Tumor Atlas Network (HTAN: https://humantumoratlas.org). We thank the scientific teams from HTAPP and HTAN for helpful discussions. We thank Soledad Villar for advice on devising and implementing the optimization routine used in this study. We thank Ania Hupalowska and Leslie Gaffney for assistance with the illustrations. We thank Alex Bloemendal for helpful discussions on the construction of the sampling models in this work.

E. B. was supported by the National Science Foundation Graduate Research Fellowship Program under Grant No. 1745302. D.S. is supported by the German Federal Ministry of Education and Research (BMBF 01ZZ2004) and was funded by an Early Postdoc Mobility fellowship (no. P2ZHP3_181475) from the Swiss National Science Foundation and was a Damon Runyon Fellow supported by the Damon Runyon Cancer Research Foundation (DRQ-03-20). B. D. was supported by the National Science Foundation (NSF) under grant DMS-1638352 and completed part of this work while visiting The Statistical and Applied Mathematical Sciences Institute in Durham, NC, under the kind support of the NSF grant DMS-1638521. S.V. was supported by the Knut and Alice Wallenberg Foundation, the Royal Swedish Academy of Sciences, Swedish Society for Medical Research and as a Knut and Alice Wallenberg Fellow at the Broad Institute of MIT and Harvard.

## Data availability

All data used in this study has been previously published and is available via the respective publications.

## Code availability

Code for tissue generation and power analysis is available at https://github.com/klarman-cell-observatory/PowerAnalysisForSpatialOmics.

## Contributions

D.S., S.V. and A.R. designed the study. E.B. designed and E.B. and B.D. implemented the tissue generation framework with supervision from D.S., S.V., and A.R.. D.S., E.B., S.V., B.D. and A.R. wrote the manuscript. All the authors read the manuscript and discussed the results.

## Supplementary figure legends

**Supplementary Fig. 1:**
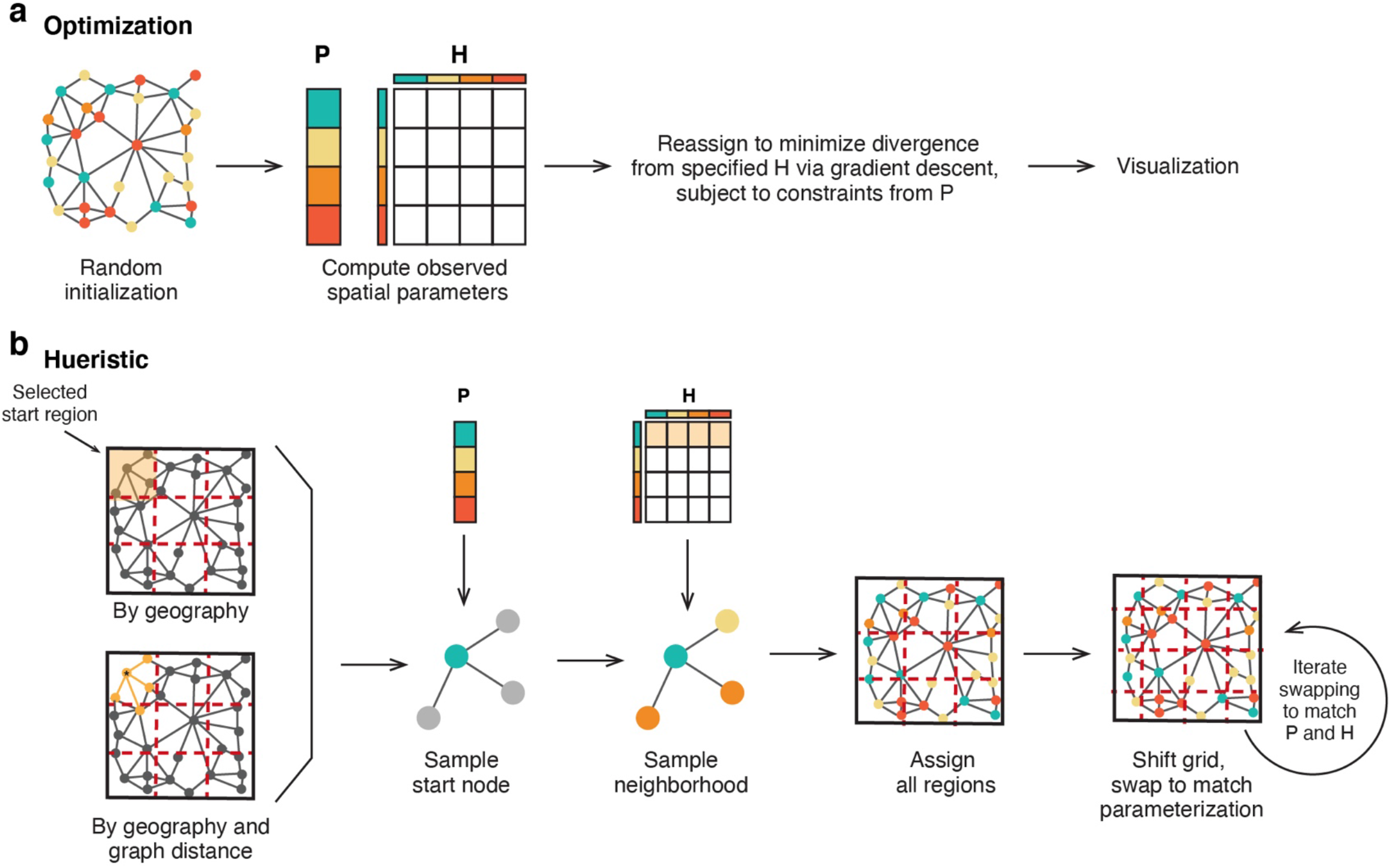
Labeling tissue scaffolds. **(a)** Overview of methods used to optimize tissue scaffold labeling. Spatial parameters (biological priors) and the tissue scaffold are provided to the assignment algorithm, which optimizes cell type labeling on the scaffold via gradient descent. **(b)** Overview of the heuristic solution used to optimize tissue scaffold labeling. Spatial parameters (biological priors) and the tissue scaffold are provided to the assignment algorithm, which attempts to generate a labeling that matches specified spatial parameters.

**Supplementary Fig. 2:**
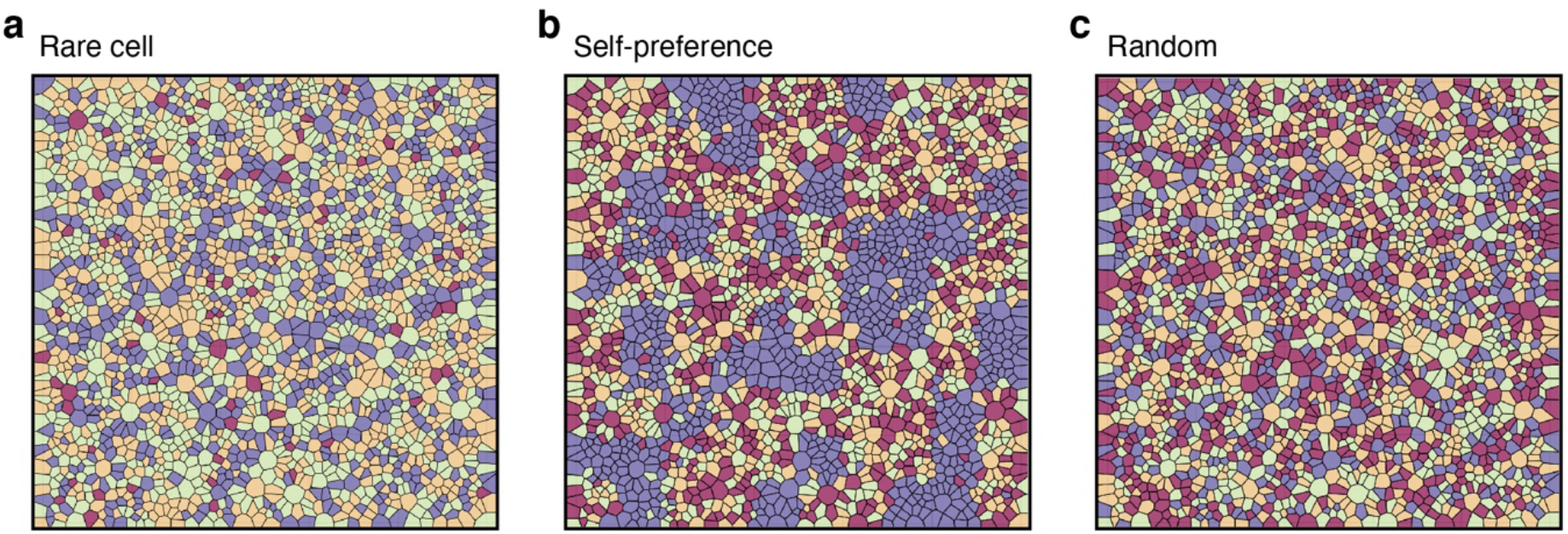
Example *in silico* tissues mimicking three biological scenarios. ISTs illustrating tissues with a rare cell type (**a,** maroon), one cell type with a strong preference to locate next to cells of the same type (**b**, purple), and no spatial constraints (**c**). Other cell types (colors) are of roughly equal abundance and have random distribution in space.

**Supplementary Fig. 3:**
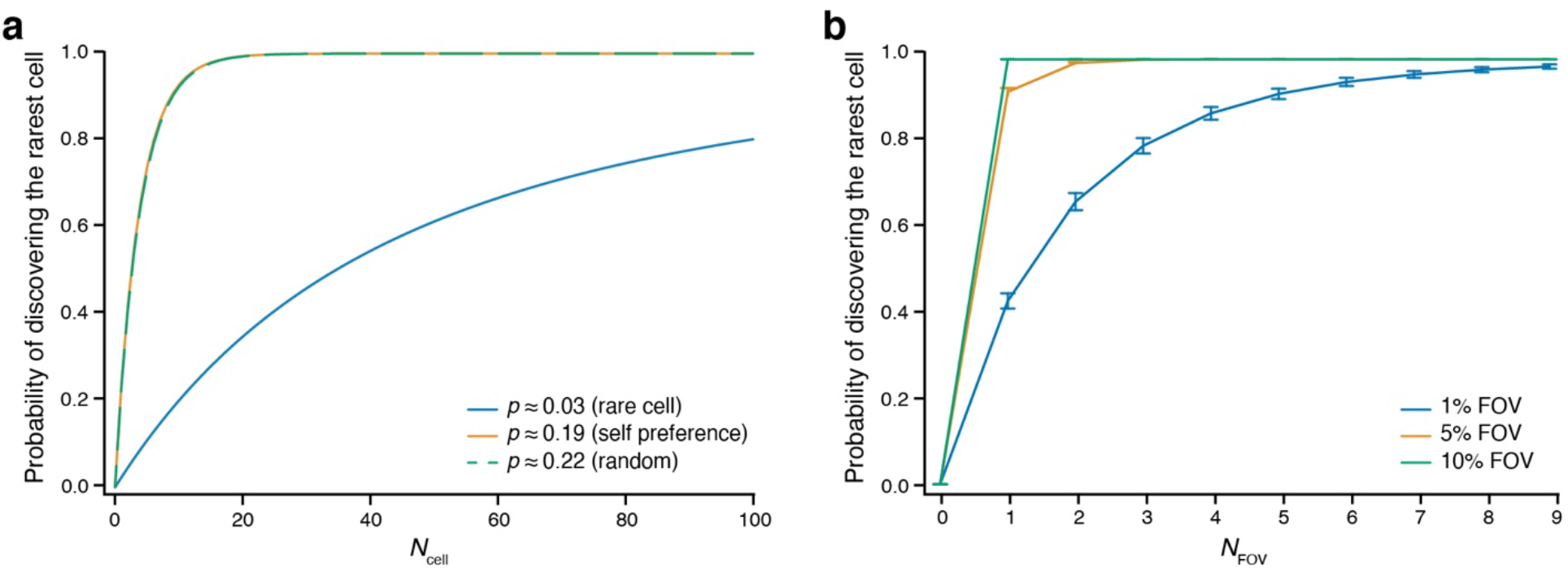
Effect of cell type abundance and FOV size on cell type recovery in small *in silico* tissues. **(a)** Impact of number of sampled cells on rare cell type detection. Beta-binomial probability (y axis) that the rarest cell type is observed at least once when sampling different numbers of cells (x axis, *N_cells_*), for each of the three *in silico* tissue types in **Supplementary Fig. 2**. **(b)** Impact of size of FOV on rare cell type detection. Beta-binomial probability (y axis) that the rarest cell type is observed in at least one FOV when sampling different numbers of FOVs (x axis), for FOVs of sizes equivalent to 1% (blue), 5% (orange), or 10% (green) of total tissue size.

**Supplementary Fig 4:**
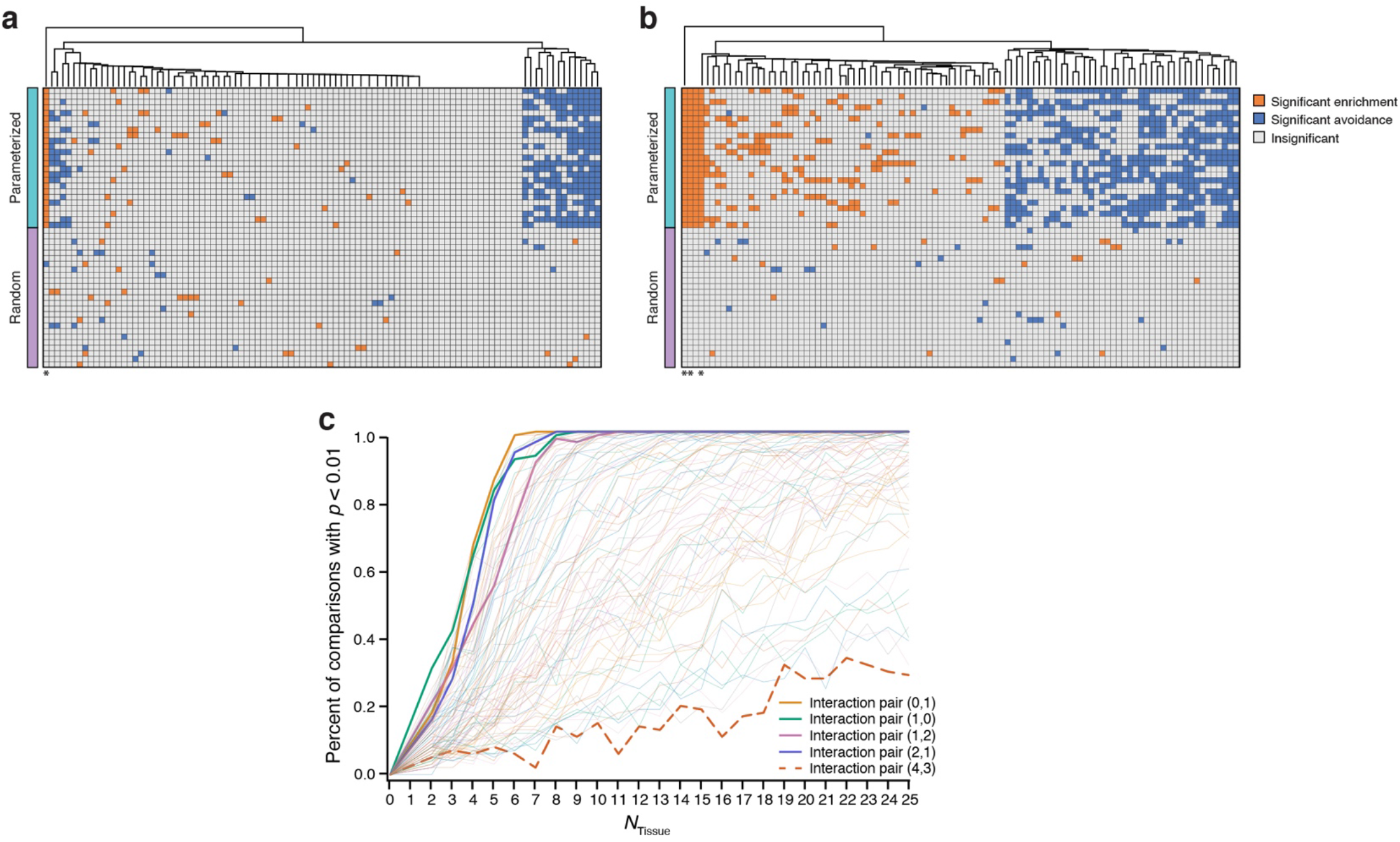
Significant pairwise interactions between cell types can be generated and detected in *in silico* data. **(a,b)** Significant pairwise interactions recovered in parametrized ISTs. Statistically significant (p<0.01, permutation test) avoidances (blue) and interactions (orange) between each pair of cell types (columns) (10 cell types, 100 pairs) in each of 50 *ISTs* (rows), 25 with random spatial pattern (spatial null, purple), and another 25 IST parametrized (cyan) with either a self-preference for one cell type (**a**) or with a significant motif of 3 interacting cell types. “*” denotes the specified significant interaction pair. (**b**). The self-preference interaction in (**a**) is in the leftmost row. “*” denotes the specified significant interaction pairs. **(c)** Impact of number of tissues on detection of significant interactions. Fraction of comparisons returning a statistically significant difference (*y* axis) between parameterized IST (with a significant 3 cell types interaction) and null ISTs (10 cell types per IST), for different numbers of ISTs per category (x axis). Interaction pairs (0,1), (1,0), (1,2), and (2,1) (highlighted traces) are the parameterized cell types representing the interaction of interest. Interaction pair (4,3) (dashed line) was not a specified relationship in the tissue generation.

**Supplementary Fig. 5:**
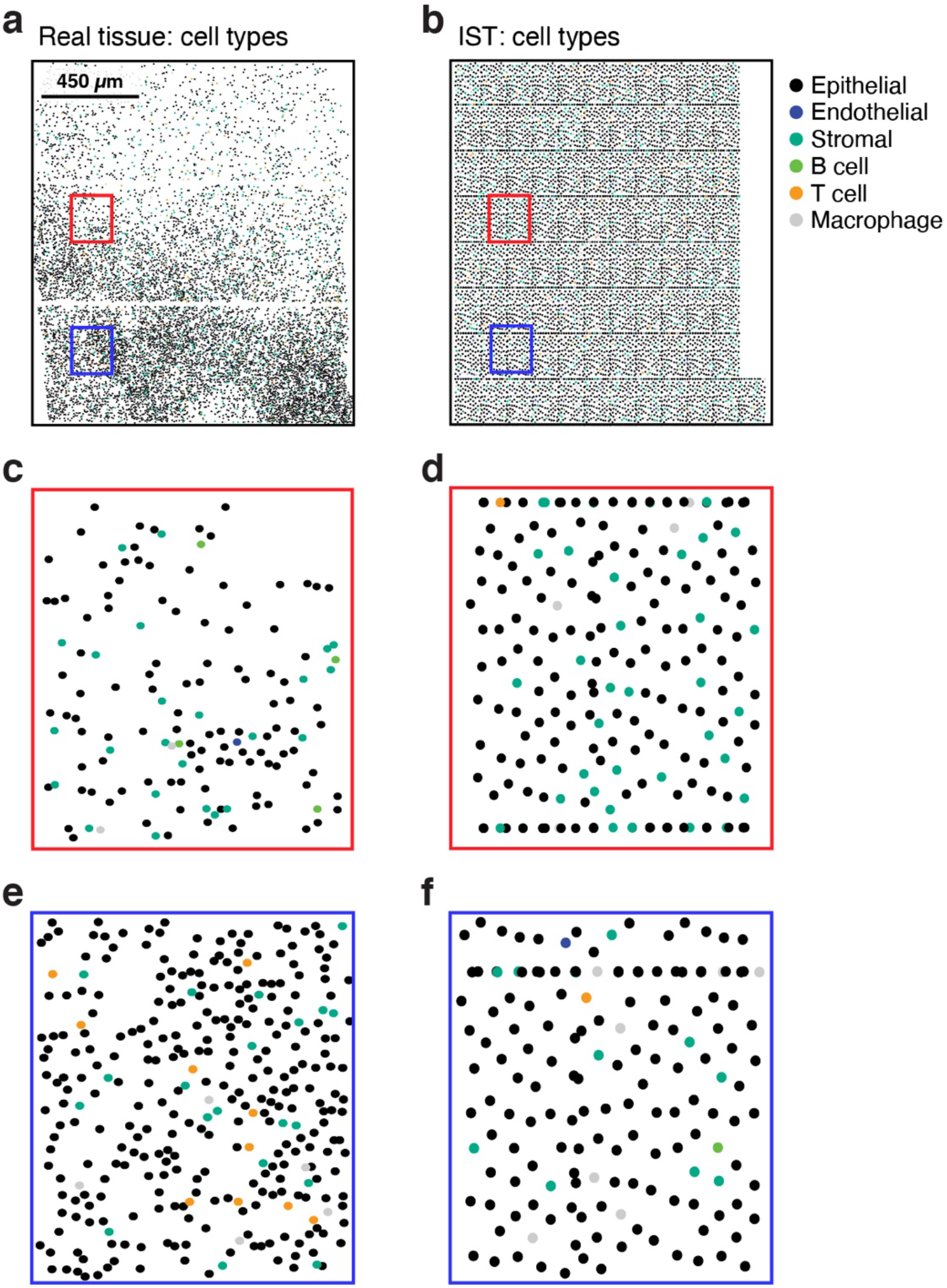
Breast cancer ISTs mimic real spatial data. **(a,b)** Parameters from real HDST data in breast cancer are used to generate an IST. Cells (dots) colored by cell type label in real HDST (**a**) and the ISR generated based on its spatial parameters (**b**). Red and blue boxes are matching regions in HDST and the IST expanded in (**c-f**). **(c-f)** Agreement in region characteristics between HDST and the generated IST. An expanded view of the red (**c,d**) or blue (**e,f**) regions in HDST (**a**) or the IST (**b**). The red region in both HDST (**c**) and IST (**d**) is dominated by epithelial cells (black) with relatively high occurrence of stromal cells (teal). The blue regions in both HDST (**e**) and IST (**f**) is also dominated by epithelial cells (black) but with a lower proportion of stromal cells (teal), and a higher abundance of T cells (orange) and macrophages (grey), as well as B cells (green) in the IST (**e**) but not HDST (**f**).

**Supplementary Fig. 6:**
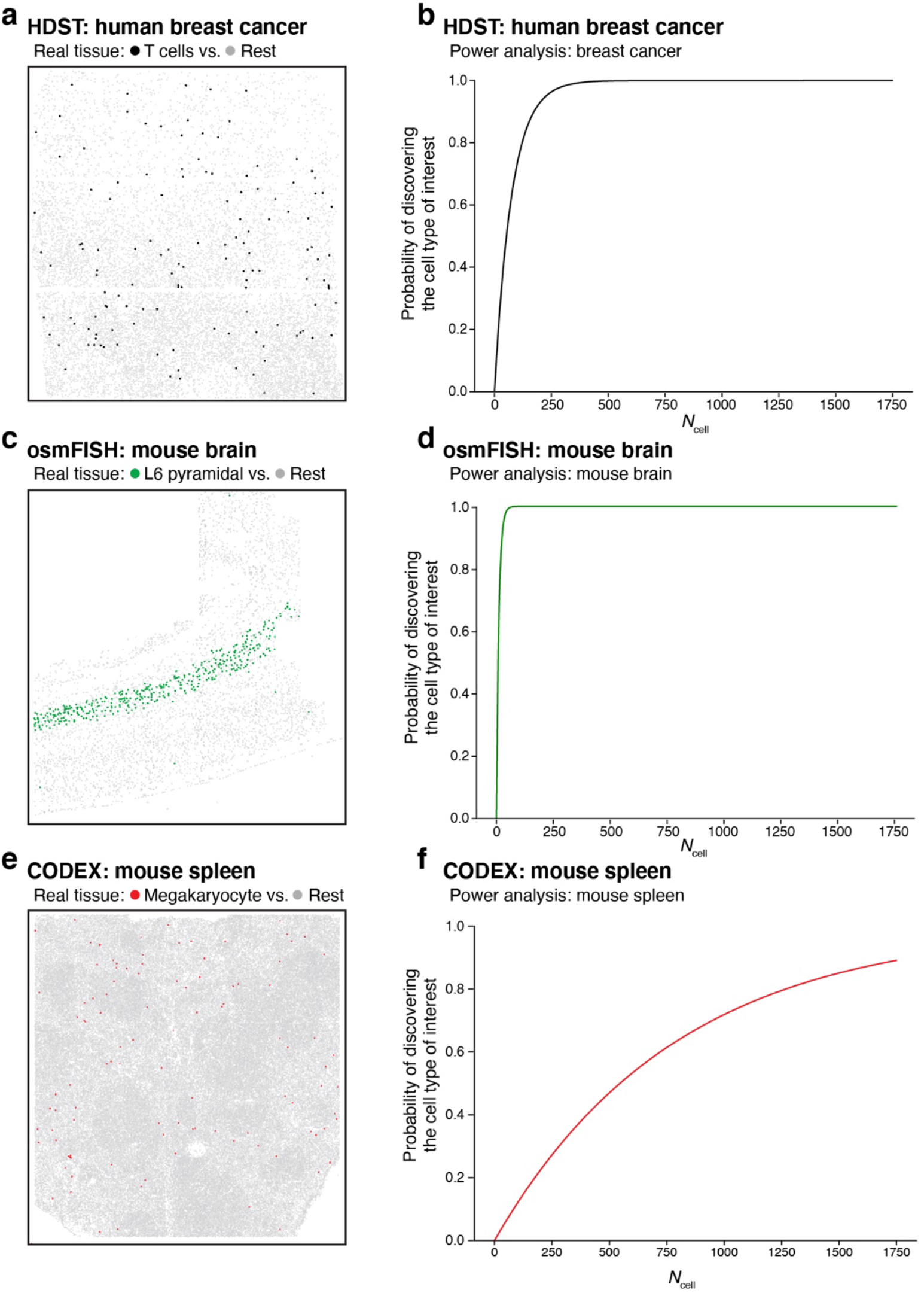
Power analysis for single cell sampling for cell type discovery. (**a,c,e**) Selected cell types for detection in three tissue types. Cell type (color) for detection in real spatial data from breast cancer (**a**, T cell), mouse cortex (**c**, L6 pyramidal neurons) and spleen (**e**, megakaryocytes) **(b,d,f)** Probability of detecting at least one cell of the type of interest (y axis) when using single cell sampling of different numbers of cells (x axis) in a single cell profiling experiment of the full tissue. Results shown for breast cancer **(b)**, mouse cortex **(d)** and spleen **(f)**data.

**Supplementary Fig. 7:**
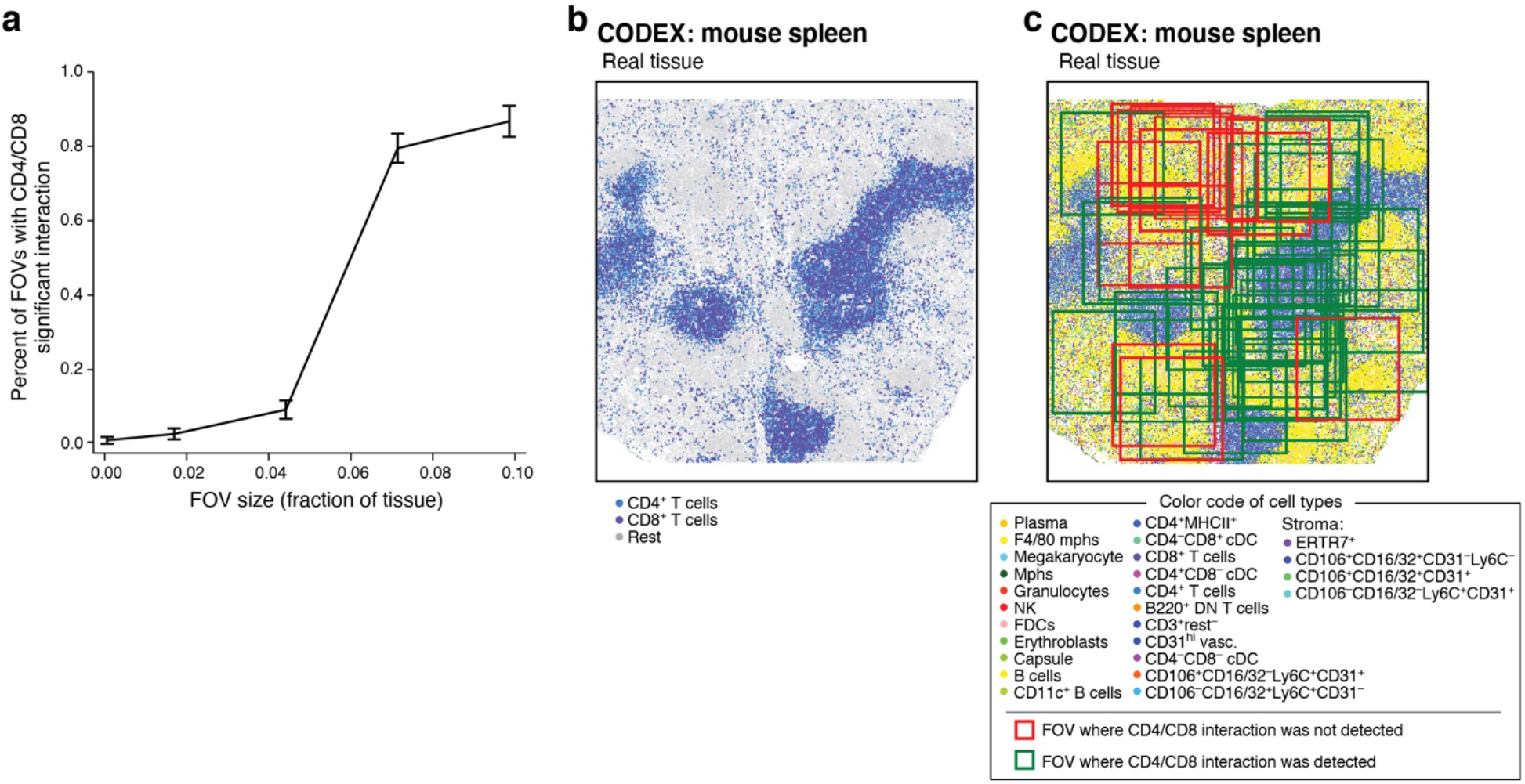
Effect of FOV size on the detection of a significant CD4^+^/CD8^+^ T cell interaction in mouse spleen. **(a)** Impact of FOV size on probability to detect cell-cell interaction. The fraction of FOVs with a significantly enriched CD4^+^/CD8^+^ T cell interaction (*y* axis) for different FOVs sizes (x axis, % of total tissue size). (**b,c**) CD4^+^ CD8^+^ T cell interactions. (**b**) CODEX data with cells (dots) colored by CD4^+^ (light blue) and CD8+ (dark blue) labels. **(c)** FOV selections (squares), each sized at 7.5% of total tissue area, where CD4^+^ CD8^+^ T cell interactions were significantly enriched (green) or not (red).

**Supplementary Fig. 8:**
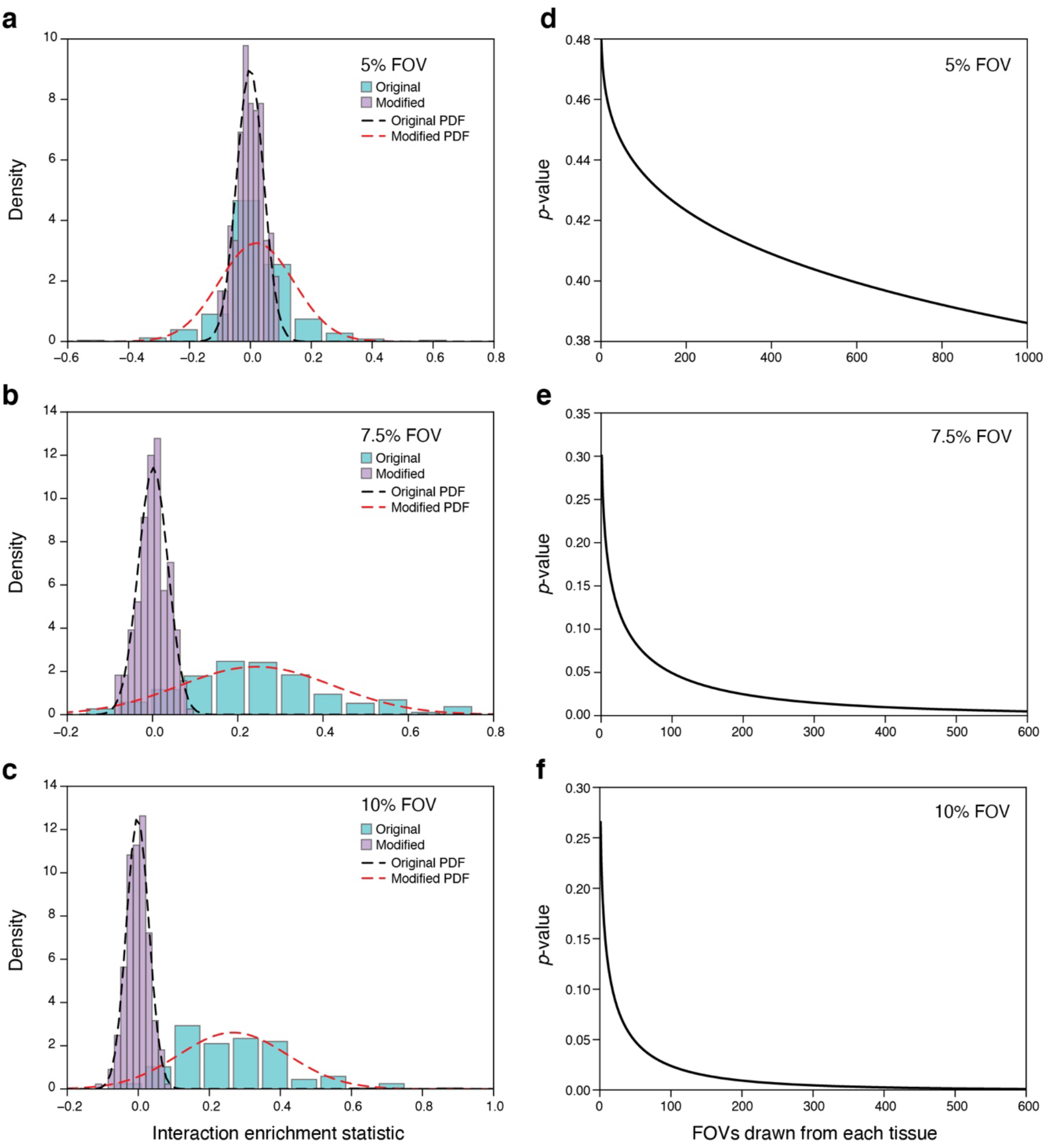
Comparison of tissue cohorts by interaction enrichment statistics (IES) **(a-c)** Larger FOVs distinguish tissues with different interaction enrichment statistics. Distribution of interaction enrichment statistics for FOVs of 5% **(a)**, 7.5% **(b)**, or 10% **(c)** absolute tissue size drawn from the original spleen tissue (cyan) or a modified dataset where the overall number of CD4^+^−CD8^+^ T cell adjacencies has been reduced by 37%, but the relative frequency of cell types and absolute structure have been preserved (magenta). Dashed lines: probability density functions for the original (red) and modified (black) tissues. **(d-f)** Power analysis for comparison of distributions of IES between original and modified tissues. Expected p-value (y axis) in a t-test comparing the IES of original and modified tissues (as in **a-c**), for different numbers of FOVs (x axis) sized at 5% (d), 7.5% (e), or 10% (f) of absolute tissue size. Smaller number of FOVs (samples) is required to distinguish the difference in interaction enrichment by the IES test (P=0.05) as the FOV size grows (~1000, ~100, and ~50, respectively).

**Supplementary Fig. 9:**
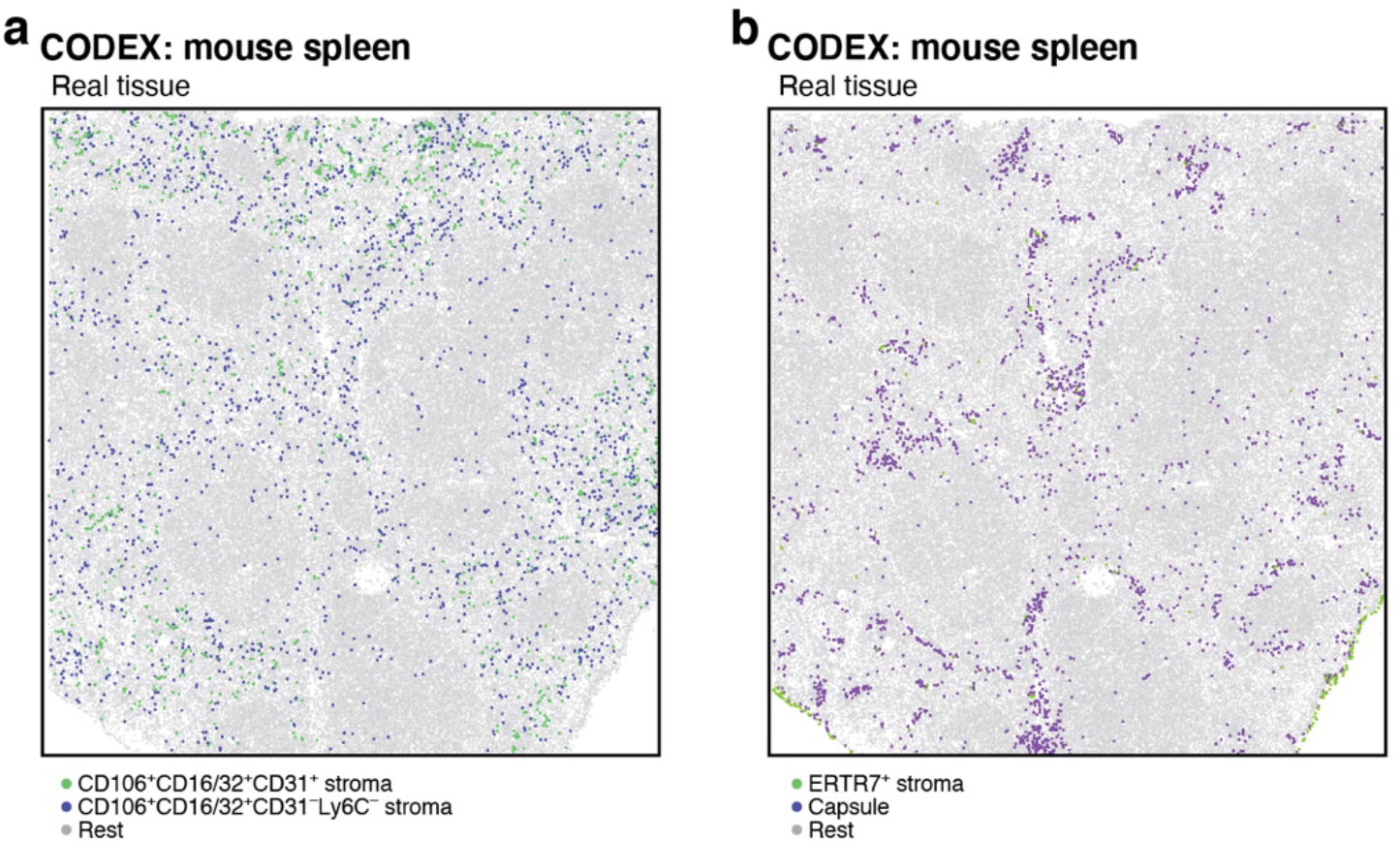
Cells participating in significant interactions that were not detected in the IST cohort are at the margins of the real tissue. Cells (dots) labeled by cell type (color) in real spleen CODEX data for a significant interaction detected by IST analysis (**a**, between CD106^+^CD16/32^+^CD31^+^ and CD106^+^CD16/32^+^CD31^-^Ly6C^-^ cells), and for a significant interaction not found in ISTs (**b**, between ERTR7^+^ cells and the capsule at the tissue edges).

