## Supplemental Table 1 for "Power analysis for spatial omics"

**Supplemental Table 1: Summary of experimental methods for spatial data generation**

| | | |
|---|---|---|
| **Transcriptomic** | *Sequencing* | Slide-seq[1] |
| | | Spatial Transcriptomics (commercially available as 10X Visium)[2] |
| | | High-definition spatial transcriptomics (HDST)[3] |
| | | *In situ* RNA sequencing[4] |
| | *Hybridization* | cyclic-ouroboros smFISH (osmFISH)[5] |
| | | sequential Fluorescence In Situ Hybridization (seqFISH)[6] |
| | | multiplexed error-robust FISH (merFISH)[7] |
| **Proteomic** | *Imaging* | CO-Detection by indEXing (CODEX)[8] |
| | | Cyclic Immunofluorescence (CyCIF) [9,10] |
| | | Multiplexed immunohistochemical consecutive staining on single slide (MICSSS)[11] |
| | *Mass spectrometry* | Imaging Mass Cytometry (IMC)[12] |
| | | Multiplexed Ion Beam Imaging (MIBI)[13] |
